# IDBSpred: An intrinsically disordered binding site predictor using machine learning and protein language model

**DOI:** 10.64898/2026.03.27.714773

**Authors:** Drew Jones, Yinghao Wu

## Abstract

Intrinsically disordered proteins (IDPs) mediate many cellular functions through interactions with structured protein partners, but predicting the corresponding binding sites on the structured partner remains challenging. Here, we present IDBSpred, a sequence-based method for residue-level prediction of IDP-binding sites on structured proteins. Training and test data were collected from the DIBS database, which contains more than 700 non-redundant IDP–protein complexes. Residue-level embeddings of structured partner sequences were generated using the ESM-2 protein language model and used as input to a multilayer perceptron classifier for binary prediction of binding versus non-binding residues. Analysis of amino acid composition showed that IDP-binding sites are enriched in aromatic residues, especially Trp, Tyr, and Phe, as well as several charged and polar residues, whereas Ala and several small or conformationally restrictive residues are depleted. The classifier achieved an ROC AUC of 0.87 and an average precision of 0.61. Structural case studies further showed that the predicted sites largely recapitulate the major experimentally defined binding interfaces. These results demonstrate that protein language model embeddings plus machine learning algorithms can effectively capture sequence features associated with IDP recognition on structured proteins. IDBSpred provides a practical framework for studying IDP-mediated interfaces and identifying potential therapeutic hotspots.

## Introduction

At least 33% of eukaryotic proteins contain regions that does not possess any stable structure in solution in their physiological condition [1]. These regions are known as intrinsically disordered regions (IDRs), and proteins that contain IDRs are called intrinsically disordered proteins (IDPs) [2]. As accumulated evidences suggesting that IDPs/IDRs are involved in carrying out versatile cellular functions [3], a new ‘disorder–function paradigm’ was proposed, in which proteins lacking a single stable conformation while alone can adopt relatively ordered structure upon encountering their binding partners [4]. This conformational plasticity leads to the fact that a single IDR can bind to several structurally diverse partners; vice versa, many different IDPs can bind to a single globular receptor [5]. As a result, many IDPs and IDRs act as “hub” molecules in the protein-protein interactions (PPIs) network [6]. These interactions dominate the functions of IDPs in many important cellular processes. Mutations of these proteins that disrupt the interactions with their binding partners can lead to various human diseases, including diabetes [7], cancer [8], and amyloidosis [9]. Therefore, IDPs have emerged as one of the prime targets for drug discovery or repurposing.

It is currently a big challenge to systematically detect interactions between IDPs and their binding partners by high-throughput methods, such as yeast-two-hybrid [10], due to the transient nature of their binding kinetics [11]. Moreover, it is even harder to characterize the structure of interactions between IDPs and their binding partners by the traditional techniques including X-ray crystallography and Cryo-electron microscopy (cryo-EM) [12]. Comparing to the time-consuming and labor-intensive experimental approaches, computational methods serve as an ideal alternative to testing conditions that are currently inaccessible in the laboratory. The applications of machine learning (ML), especially deep learning such as AlphaFold [13], to study PPIs has gained enormous attentions [14, 15]. Unfortunately, these methods were mainly trained on large-scale databases of folded proteins, and are thus much more sensitive to recognizing the specific interactions between proteins with well-defined tertiary structures. Whether these methods are able to model the fuzzy interactions of IDPs is still an unanswered question. Therefore, the development of new computational methods that specifically focus on the interactions between IDPs and their binding partners is highly demanding.

Most existing computational approaches have concentrated on detecting binding-prone segments within intrinsically disordered sequences, rather than identifying the corresponding residues on their folded partners. Representative methods include ANCHOR, which infers disordered binding regions from sequence-derived energetic considerations [16], and later machine-learning predictors such as MoRFpred [17-20] and DISOPRED3 [21, 22], which use sequence-derived features to identify disorder-mediated protein-binding regions. However, the reciprocal problem, predicting which residues in the structured partner mediate binding to an IDP, has received much less attention, although it is extremely important in terms of revealing targetable interaction hotspots for designing peptide-based therapeutics and other drugs that modulate disease-relevant IDP-mediated protein–protein interactions. A recent advance in this direction is Disobind, which uses protein language model embeddings to predict partner-dependent contact maps and interface residues for IDR–partner interactions from sequence [23]. Nevertheless, computational methods specifically designed for residue-level identification of IDP-binding sites on structured partner proteins are still scarce, motivating the development of new approaches for this task.

In this work, we present IDBSpred (**Figure 1**), a computational framework for residue-level prediction of IDP-binding sites on structured partner proteins. IDBSpred was developed using more than 700 non-redundant IDP–protein complexes collected from the DIBS database [24], in which residues on the structured binding partner were annotated as either IDP-binding or non-binding. To capture informative sequence context, we extracted residue-level embeddings from the ESM-2 protein language model [25] and used these representations as input to a multilayer perceptron classifier for binary prediction. Using this strategy, IDBSpred achieves strong overall performance in distinguishing binding from non-binding residues and is able to recover the major interface regions in structured binding partners of IDP proteins. By focusing specifically on the structured side of IDP-mediated interactions, IDBSpred provides a practical approach for identifying potential IDP-recognition surfaces and offers a useful tool for studying the molecular basis of IDP binding and for guiding the design of therapeutics targeting IDP-mediated protein–protein interactions.

**Figure 1:**
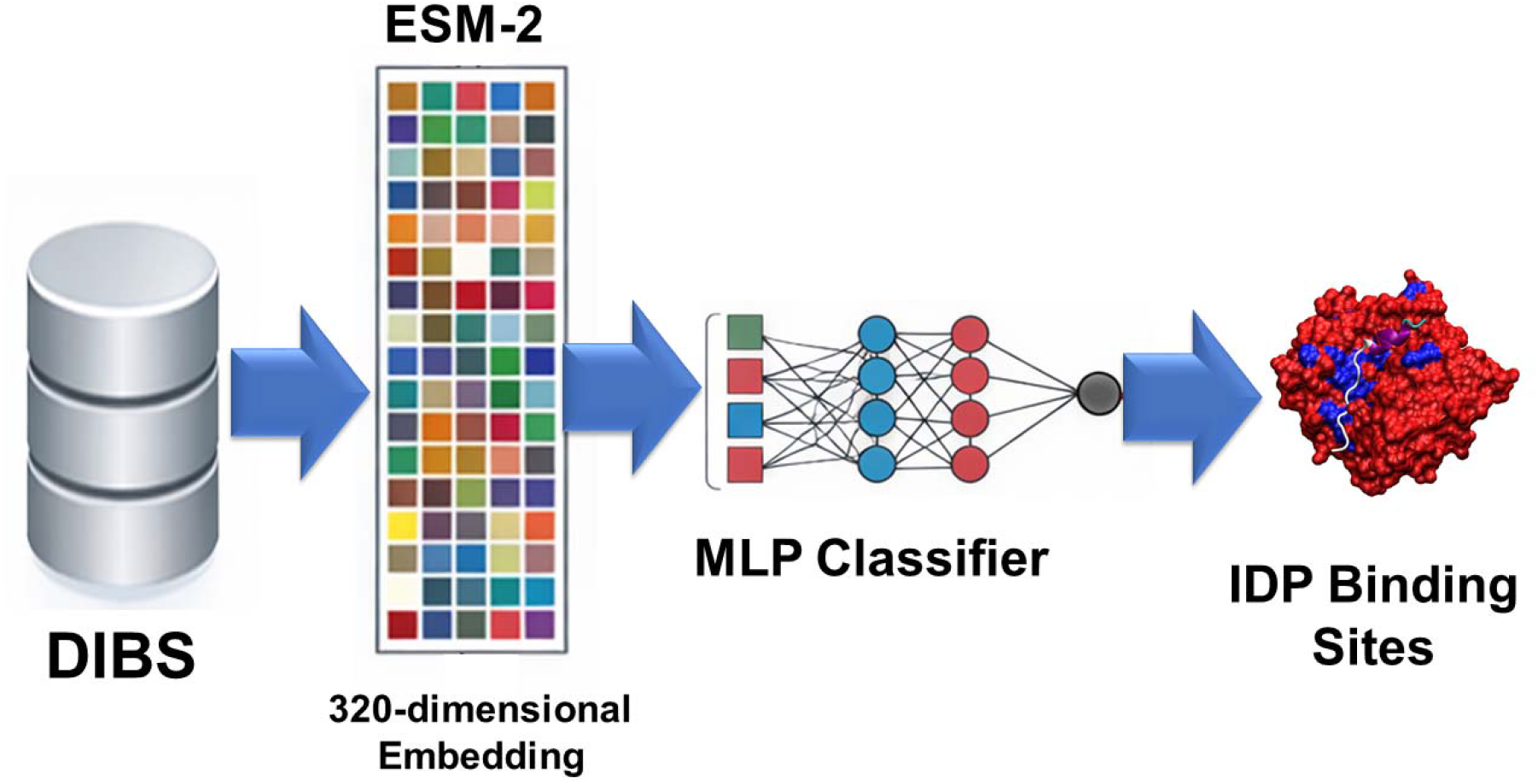
Overview of the IDBSpred workflow. Schematic illustration of the IDBSpred framework for residue-level prediction of IDP-binding sites on structured partner proteins. IDP–protein complexes were collected from the DIBS database, and residues in the structured partners were annotated as either IDP-binding or non-binding. Residue-level sequence embeddings were generated using the ESM-2 protein language model, producing 320-dimensional feature vectors for each residue. These embeddings were then used to train a multilayer perceptron (MLP) classifier to predict whether a residue is part of an IDP-binding site. The dataset was divided into training and testing sets for model development and evaluation.

## Model and Methods

### Based Dataset construction

Training and evaluation data were collected from the DIBS database [24], which contains more than 700 non-redundant interacting complexes formed between intrinsically disordered proteins (IDPs) and their structured binding partners. For each complex, the structured partner protein sequence was used for residue-level feature extraction and binding-site annotation.

Residues in the structured partner were labeled according to whether they participate in binding to the IDP. Residues that directly interact with the IDP were defined as positive samples (IDP-binding residues), whereas residues in the same structured proteins that do not interact with the IDP were defined as negative samples (non-IDP-binding residues). Thus, the task was formulated as a binary classification problem at the residue level.

### Protein language model embeddings

To represent each residue numerically, we used the ESM-2 protein language model [25] to generate sequence embeddings for all structured binding partners in the DIBS dataset. For each residue, a 320-dimensional embedding vector was extracted from the model and used as the input feature for classification. These embeddings were expected to capture contextual sequence information relevant to residue function and binding propensity.

### Neural network classifier

A multilayer perceptron (MLP) was used to classify each residue as either an IDP-binding site or a non-binding residue. The model took a 320-dimensional residue embedding as input and produced a single binary output score. The architecture consisted of one fully connected hidden layer with 128 neurons, followed by a ReLU activation function and a dropout layer with a dropout rate of 0.3. The final output layer mapped the hidden representation to a single logit value corresponding to the predicted binding probability.

### Model training

All positive and negative residues extracted from the database were randomly divided into training and test sets. A total of 80% of residues were used for training, and the remaining 20% were reserved for testing. The binary labels and corresponding residue embeddings were organized into tensor datasets and loaded in mini-batches for model training and evaluation.

The MLP classifier was implemented in PyTorch [26]. Model parameters were optimized using the Adam optimizer with a learning rate of 1×10^-3^. Binary classification loss was calculated using binary cross-entropy with logits. Training was performed for 25 epochs with a batch size of 32. During each epoch, the model was trained on the training set and then evaluated on the held-out test set to monitor validation loss.

### Code availability

All relevant source codes of the classifier can be found in the GitHub repository: https://github.com/wulab-github/IDBSpred

## Results

We first investigated the preference of all 20 different amino acids to be presented at the IDP binding sites. **Figure 2** shows the amino acid composition bias of IDP-binding sites in all structured partner proteins in the DIBS database, measured as the natural logarithm of the ratio between the frequency of each amino acid at IDP-binding sites and its overall frequency in proteins. Positive values indicate enrichment at IDP-binding sites, whereas negative values indicate depletion. Among the 20 amino acids, Trp shows the strongest enrichment, followed by Tyr, suggesting that aromatic residues are highly favored at interfaces that recognize IDPs. Phe is also enriched, further supporting the importance of aromatic side chains in mediating IDP recognition. In addition, positively charged and polar residues such as Arg, His, Lys, Met, and Asn show moderate enrichment, indicating that electrostatic interactions and hydrogen bonding may also contribute substantially to IDP binding. In contrast, residues such as Ala are strongly depleted, and Pro, Ser, Gly, Cys, Glu, Asp, and Val also occur less frequently than expected at IDP-binding sites. Overall, these results suggest that IDP-binding interfaces on structured proteins preferentially contain aromatic, charged, and interaction-capable residues, while small aliphatic or conformationally restrictive residues are less favored. This pattern is consistent with the idea that recognition of disordered partners relies on a combination of hydrophobic packing, aromatic contacts, and flexible polar interactions rather than nonspecific surface exposure alone.

**Figure 2.**
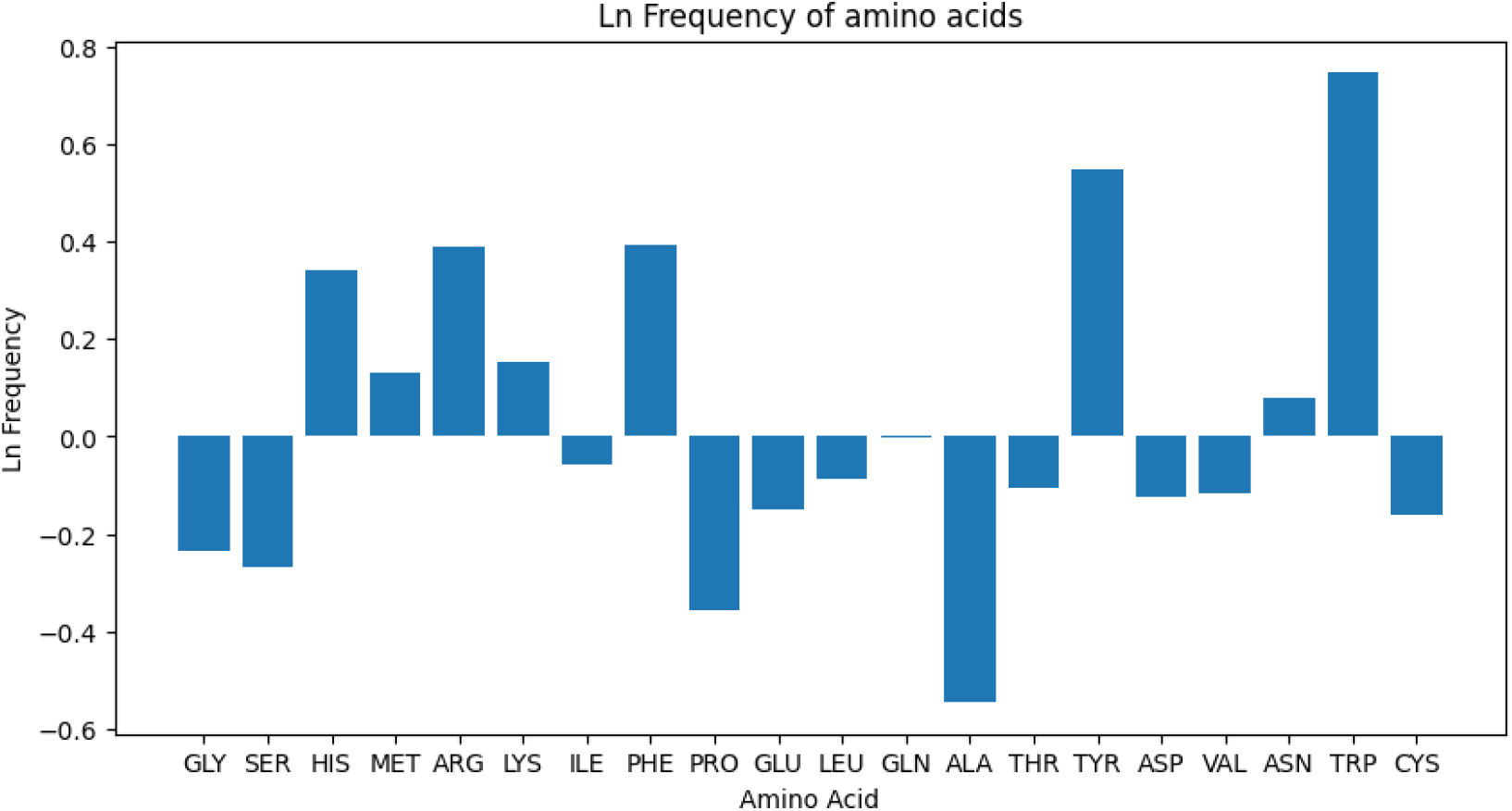
Amino acid preferences at IDP-binding sites in structured partner proteins. Bar plot showing the residue composition bias of IDP-binding sites in structured binding partners from the DIBS database. Values represent the natural logarithm of the ratio between the frequency of each amino acid at IDP-binding sites and its overall background frequency in proteins. Positive values indicate enrichment at IDP-binding sites, whereas negative values indicate depletion. Aromatic residues, especially Trp and Tyr, are strongly enriched, while residues such as Ala and Pro are underrepresented.

**Figure 3** presents the overall performance of the MLP classifier for identifying IDP-binding residues in structured partner proteins from residue-level ESM-2 embeddings. As shown in **Figure 3a**, the training loss decreases steadily over the course of 25 epochs, indicating that the model progressively captures informative patterns from the embedding features. The validation loss also declines markedly during the early epochs and then remains relatively stable with only modest variation, suggesting that the model reaches a consistent level of generalization on the held-out test data. These learning curves indicate that the ESM-2 residue representations contain sufficient information to support classification of IDP-binding versus non-binding residues using a relatively simple neural network architecture.

**Figure 3.**
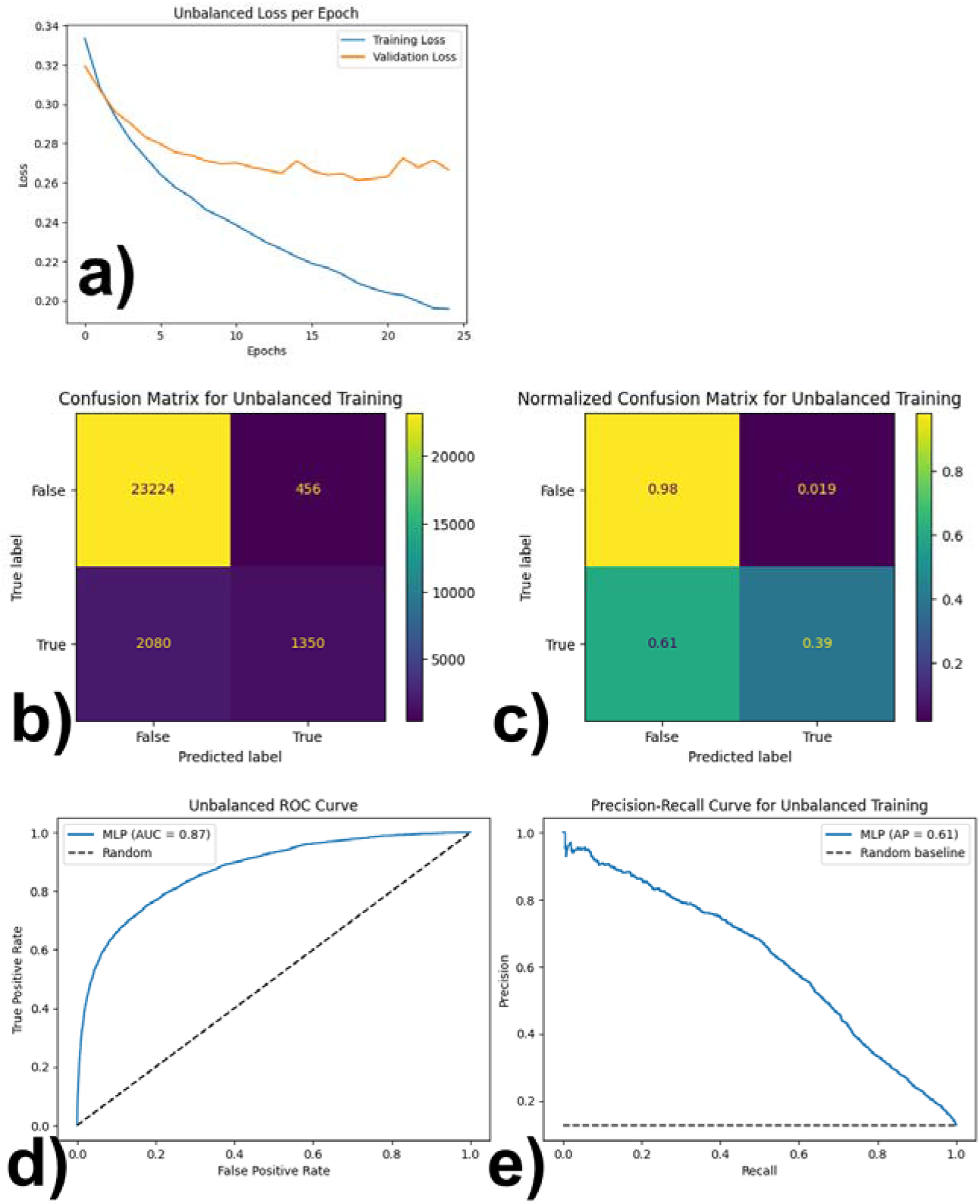
Performance evaluation of IDBSpred on residue-level prediction of IDP-binding sites. **(a)** Training and validation loss curves over 25 epochs. **(b)** Confusion matrix for classification of IDP-binding and non-binding residues on the test set. **(c)** Normalized confusion matrix showing class-wise prediction performance. **(d)** Receiver operating characteristic (ROC) curve, with an area under the curve (AUC) of 0.87. **(e)** Precision-recall (PR) curve, with an average precision of 0.61. Together, these results show that IDBSpred effectively distinguishes IDP-binding residues from non-binding residues.

The confusion matrix in Figure 3b further illustrates the classification behavior of the model. A large number of non-binding residues are correctly assigned as negatives, whereas a smaller but substantial fraction of true binding residues are correctly identified as positives. The normalized confusion matrix in **Figure 3c** makes this trend clearer by showing that the classifier achieves very high accuracy for the negative class, while its sensitivity for the positive class is lower. This result indicates that the model is particularly effective at distinguishing non-binding residues from the background surface, but still misses a portion of true IDP-binding residues. Given that residue-level binding-site prediction is inherently imbalanced, with far fewer binding residues than non-binding residues, this pattern is not unexpected and reflects the greater difficulty of detecting the minority interface class.

Despite this asymmetry between classes, the model shows strong overall discrimination ability. The ROC curve in **Figure 3d** gives an AUC of 0.87, indicating that the classifier can separate binding from non-binding residues with good reliability across different decision thresholds. Similarly, the precision–recall curve in Figure 3e yields an average precision of 0.61, demonstrating that the predictor retains substantial utility when evaluated specifically on the positive class, which is the more challenging and biologically relevant target. Taken together, these results show that the combination of ESM-2 embeddings and a simple MLP provides an effective framework for residue-level prediction of IDP-binding sites on structured proteins, with especially strong performance in identifying non-binding residues and encouraging overall ability to recover true interface residues.

**Figure 4** compares experimentally defined IDP-binding sites with the sites predicted by the model for three representative complexes: 2MZD [27], 4GF3 [28], and 4L67 [29]. In each row, the left panel shows the real binding sites and the right panel shows the predicted binding sites on the structured partner surface. Blue marks binding-site residues, while red marks non-binding residues. Overall, the figure shows that the model captures the main spatial location and shape of the binding interface in all three examples. For 2MZD in **Figure 4a**, the predicted interface largely overlaps with the true interface and correctly identifies the continuous groove contacted by the IDP. The prediction reproduces the major interface patch, although some local over-prediction is visible around the edge of the surface.

**Figure 4.**
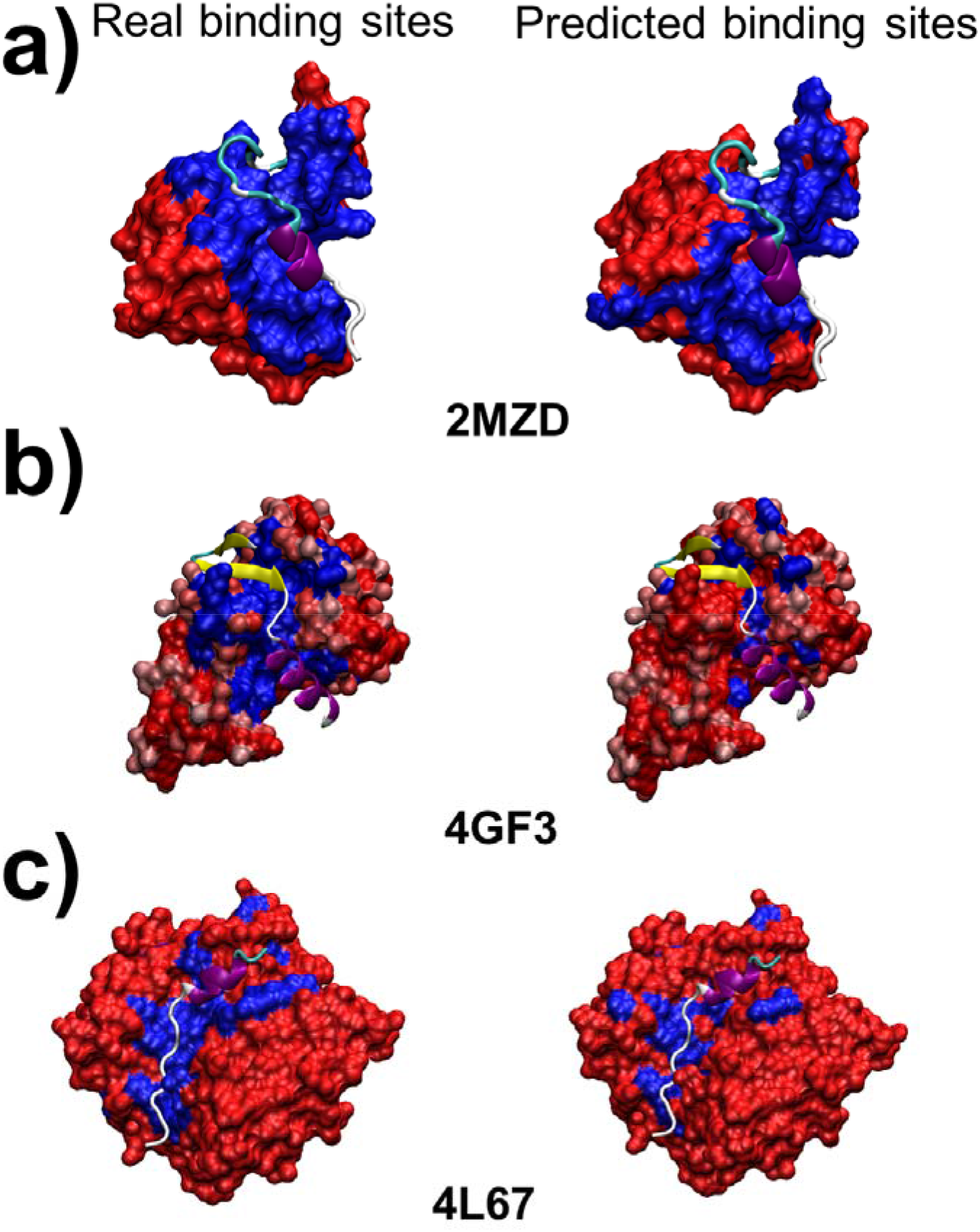
Representative examples of predicted IDP-binding sites on structured partner proteins. Comparison between experimentally annotated and predicted IDP-binding sites for three representative complexes: **(a)** 2MZD, **(b)** 4GF3, and **(c)** 4L67. In each case, the left panel shows the experimentally defined binding sites and the right panel shows the sites predicted by IDBSpred. The IDP proteins are shown in cartoon representation and their structured binding partners are shown in surface representation. Blue indicates IDP-binding residues, and red indicates non-binding residues. The model successfully recovers the major interface regions in all three examples, although some discrepancies remain at interface boundaries.

For 4GF3 in **Figure 4b**, the model again recovers the dominant binding region along the central surface contacted by the disordered partner. Compared with the native annotation, the predicted sites are somewhat more fragmented and include several extra patches, suggesting a tendency toward false-positive expansion on nearby surface residues. Even so, the core interaction region is still correctly localized. For 4L67 in **Figure 4c**, the binding interface is smaller and more localized. The model identifies the principal binding groove and preserves the overall placement of the interface, but the predicted patch is narrower than the real one in some regions and misses part of the full experimental surface. This indicates that the method can detect the correct binding neighborhood even when the interface is more compact, though some false negatives remain.

Taken together, these examples suggest that the model is effective at learning the interface propensity landscape of IDP-binding partners from residue-level PLM embeddings. The predictions are strongest at identifying the core binding region, while errors mainly appear at the interface boundary, where distinguishing true contacting residues from nearby exposed residues is more difficult. This pattern is consistent with a model that captures the global geometry of the binding surface but still has limited precision for exact residue-level delineation.

## Conclusions

In this study, we developed IDBSpred, a residue-level predictor for identifying IDP-binding sites on structured partner proteins. Using more than 700 non-redundant IDP–protein complexes from the DIBS database, we constructed a binary classification framework in which residues on the structured partner were labeled as either IDP-binding or non-binding. Residue representations extracted from the ESM-2 protein language model were used as input features for a simple multilayer perceptron classifier. Despite the simplicity of the architecture, the model achieved strong discrimination between binding and non-binding residues, with an ROC AUC of 0.87 and an average precision of 0.61. Structural case studies further showed that the predicted sites largely overlap with experimentally defined interfaces and can successfully recover the major binding regions in representative complexes.

Our analysis of amino acid composition revealed that IDP-binding sites on structured proteins are enriched in aromatic, charged, and polar residues, suggesting that these interfaces possess characteristic physicochemical features that distinguish them from general protein surfaces. These findings support the idea that IDP recognition is not random, but is mediated by specific residue environments that can be captured by protein language model embeddings. Together, these results demonstrate that sequence-derived embeddings from large protein language models contain sufficient information to enable effective prediction of IDP-binding sites on structured proteins.

Although the current model shows encouraging performance, especially in identifying the major interface regions, prediction of the minority positive class remains more challenging than prediction of non-binding residues. Future improvements may be achieved by incorporating structural context, surface accessibility, evolutionary conservation, or partner-aware information into the framework. Nevertheless, IDBSpred provides a useful first step toward systematic characterization of IDP-mediated interfaces and offers a practical computational tool for studying IDP interactions and identifying potential therapeutic hotspots for peptide- or small-molecule-based intervention.

## Acknowledgement

This work was supported by the National Institutes of Health under grant number R01GM120238, the United States–Israel Binational Science Foundation Project Number: 2023336, and the Einstein 2030 Seed Fund. The work was also partially supported by a start-up grant from Albert Einstein College of Medicine.

## Author Contributions

D.J. and Y.W. designed research; D. J. and Y.W. performed research; D.J., and Y.W. analyzed data; D.J. and Y.W. wrote the paper.

